# AmPair: automating housekeeping-gene primer design for species-level metataxonomics

**DOI:** 10.64898/2026.08.25.746527

**Authors:** Xinming Xu, Xinbin Yang

## Abstract

Amplicon sequencing of the 16S rRNA gene is the most widely used approach for profiling bacterial communities, but its taxonomic resolution is typically limited to the genus level. Many species carry multiple divergent 16S rRNA alleles that overlap across species boundaries, an ambiguity that even full-length, long-read sequencing cannot fully resolve. Shotgun metagenomics achieves species-level resolution but remains costly, particularly when only a single genus is of interest. Amplicon sequencing of rapidly evolving, protein-coding housekeeping genes offers a cost-effective alternative, yet no tool exists to identify suitable primer sets for a given target taxon. Here we present AmPair, a Snakemake pipeline that, given a target genus and one or more candidate housekeeping genes, designs and ranks primer pairs binding conserved regions while flanking a variable region capable of species-level discrimination, and validates them *in silico* across all available genomes. Using the genus *Bacillus* and the housekeeping gene *tuf* as a case study, the primer set recommended by AmPair amplified 99% of 2,392 genomes; only 0.04% carried multiple alleles and none showed inter-species allele overlap, compared with 91.41% and 69.49%, respectively, for the standard 16S rRNA V1–V9 region. Applied to a *Bacillus* community profiled by Nanopore sequencing, the same primers resolved closely related species. AmPair thus offers a generalizable and accessible route to species-level community profiling.

## Introduction

Amplicon sequencing enables the sensitive detection of genetic variation embedded within complex genomic backgrounds [1]. It is typically restricted to target regions of 100-500 nucleotides to fit short-read high-throughput sequencing platforms [2,3]. Among such applications, profiling of the 16S rRNA gene in bacteria and the internal transcribed spacer (ITS) region in fungi has become the predominant strategy for characterizing microbial community composition [4,5]. For bacterial profiling in particular, sequencing efforts are most frequently directed at the V3–V4 hypervariable region, one of the nine variable regions of the 16S rRNA gene [6]. However, this short-read, amplicon-based approach has two notable limitations. First, it is susceptible to identification bias arising from chimeric sequences generated during the PCR amplification step of library construction. Second, taxonomic assignment based on widely used 16S rRNA gene reference databases is generally confined to the genus level. Recently, these problems appear to be circumvented by the advent of third-generation sequencing platforms, namely Oxford Nanopore Technologies (ONT) and Pacific Biosciences (PacBio) [7,8]. Nevertheless, full-length, long-read 16S rRNA amplicon sequencing still lacks sufficient resolution for certain genera [9,10]. Many bacterial species carry multiple, divergent 16S rRNA gene alleles, and the alleles present within a genus are often not distinctive enough to separate species reliably [11,12]. For instance, across 856 complete genomes of the genus *Bacillus*, spanning 47 recognized species, the genus exhibits 93.93% 16S rRNA gene allele multiplicity and 55.32% inter-species allele overlap [13,14]. As a consequence, a single *Bacillus subtilis* genome may contain (i) a 16S rRNA gene copy identical to that found in another species, (ii) multiple divergent intragenomic variants, or (iii) several variants, each shared with a different non-*subtilis* species. Comparable patterns are observed in other genera, such as *Pseudomonas* and *Staphylococcus* [15,16]. Because these taxa are of considerable medical and industrial significance, the ambiguity inherent in 16S rRNA gene amplicon data may lead to unreliable diversity estimates, a problem that even full-length long-read sequencing may not overcome.

To achieve species-level identification, one option is shotgun metagenomic sequencing, which captures all genomes in a sample and thereby resolves both taxonomic composition and functional gene potential; this approach, however, remains relatively expensive. Moreover, when only a single genus is of interest, sequencing the entire community is unnecessarily costly. An alternative is target a housekeeping gene, which offers higher discriminatory power and enables species-level identification, rather than the conventional 16S rRNA gene. Housekeeping-gene amplicon sequencing therefore provides a cost-effective means of distinguishing species within mixed communities. To date, however, no tool exists to recommend which housekeeping-gene primers should be used for a given target. We therefore developed AmPair, which allows users to identify amplicon primer sets suitable for species-level discrimination within mixed communities, thereby resolving taxonomic composition without sequencing the community in full.

## Methods Workflow

AmPair is a Snakemake pipeline, freely available at https://github.com/Xinming9606/AmPair (Figure 1). Given a target bacterial genus and one or more housekeeping genes with high discriminatory power, the workflow proceeds as follows. All complete-level assemblies of the genus are retrieved from NCBI with ncbi-genome-download (https://github.com/kblin/ncbi-genome-download). For each gene, sequences are extracted from the CDS and RNA genomic FASTA files and pooled; redundant sequences are then clustered with vsearch (cluster_fast, both strands) at 97% identity, and the resulting centroids are aligned with MUSCLE v5 [17,18].

**Figure 1.**
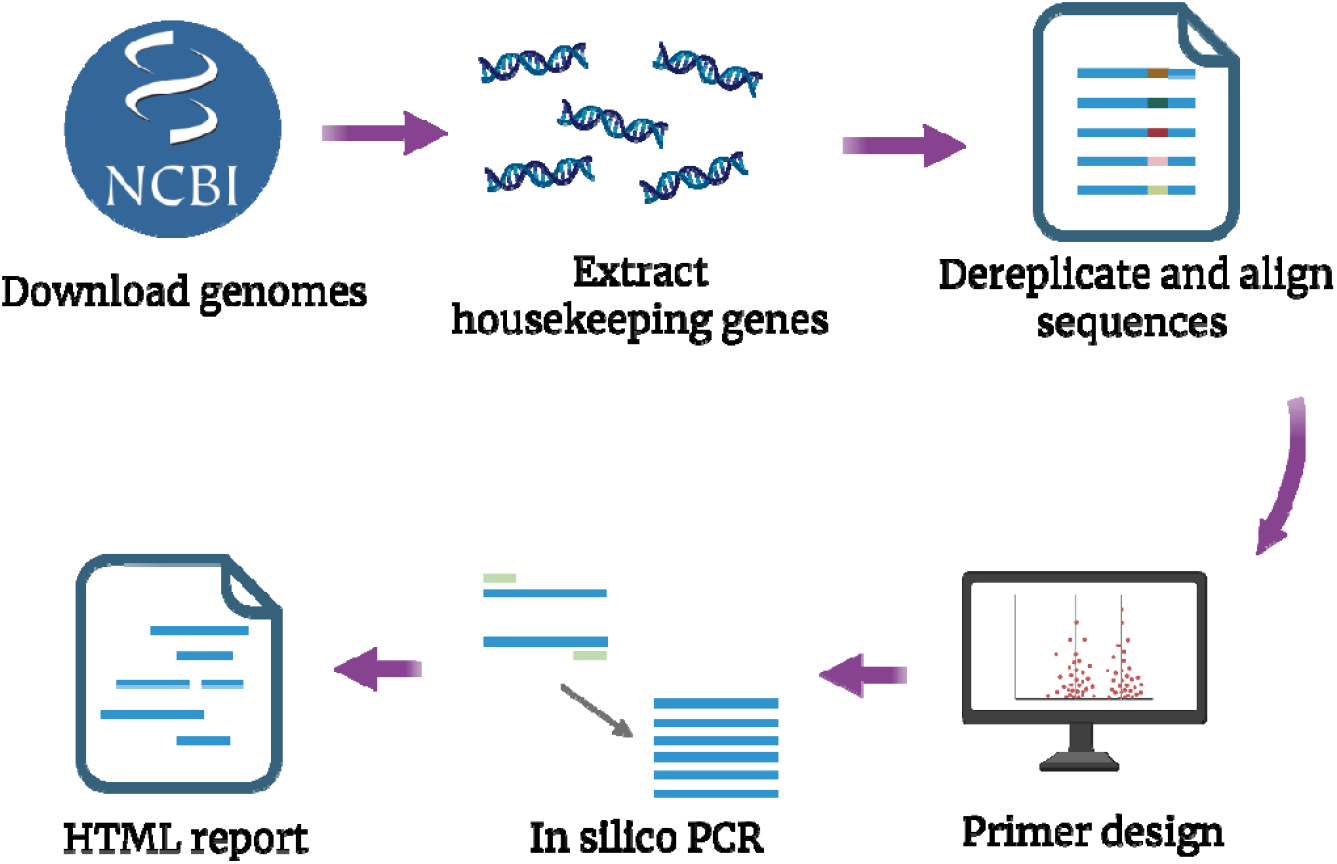
Schematic overview of the AmPair workflow. The user provides the name of a target genus, together with one or more housekeeping genes of potentially high discriminatory power. Complete assembled genomes are downloaded from NCBI, and for each housekeeping gene the corresponding sequences are extracted across all genomes, dereplicated, and aligned. Primer pairs are then designed to bind conserved regions while flanking a variable region that provides species-level discrimination. The top-ranked primer pair is validated in silico against all input genomes, and the results are compiled into per-gene and cross-gene HTML reports.

**Figure 2.**
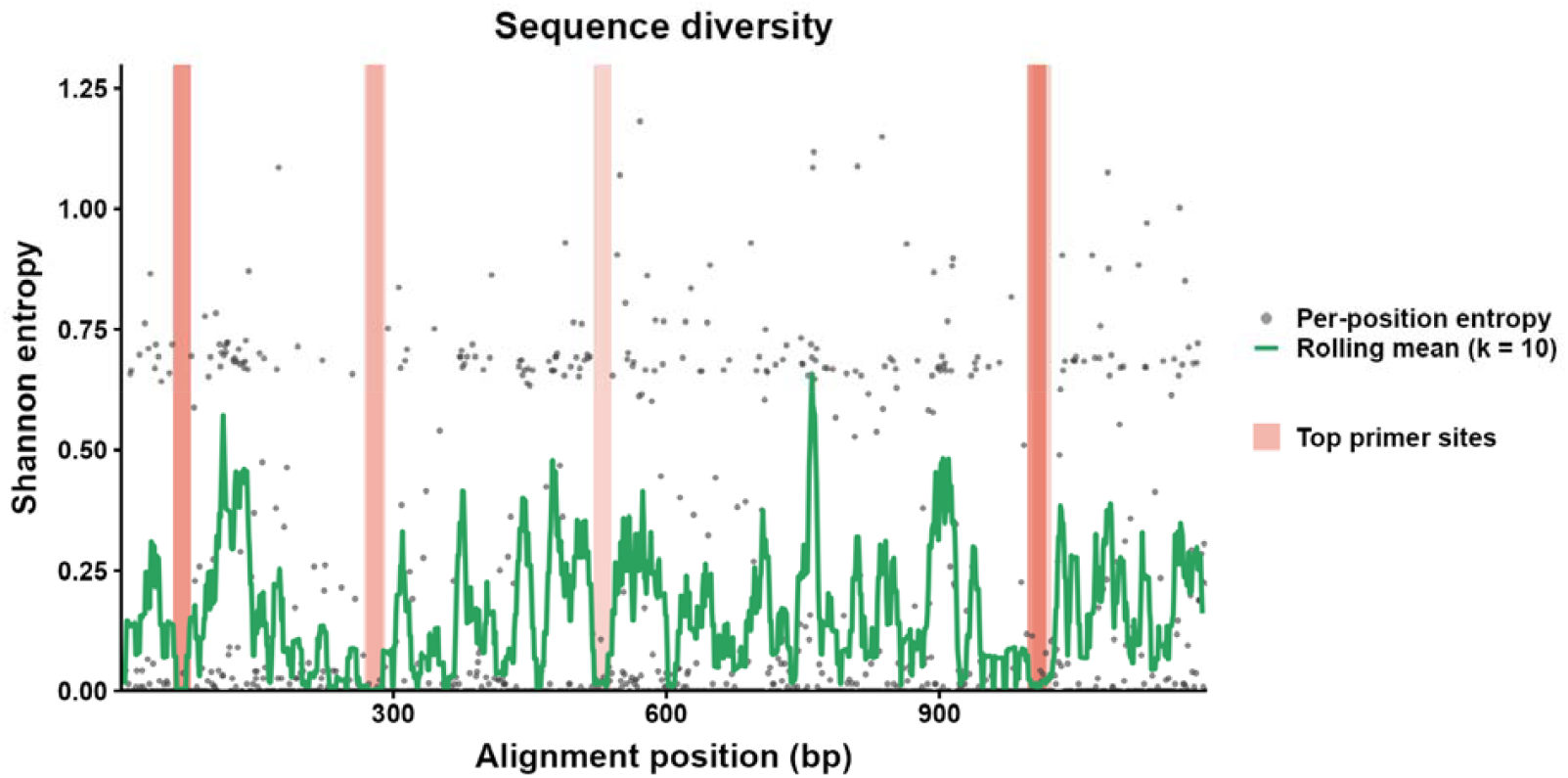
Per-position Shannon entropy across the alignment. Grey dots indicate the entropy at each position, and red rectangles mark the binding sites of the top-scoring primer pair.

Primer design follows a single principle: primers are placed in the most conserved, low-diversity regions of the alignment, whereas the intervening amplicon should be sufficiently variable to resolve taxa at the species level. For each alignment column, Shannon entropy is calculated from the observed base distribution across the aligned sequences. A majority-rule consensus is derived separately and is used only to provide the consensus and IUPAC representation from which primer windows are taken; entropy is not calculated from the consensus sequence. Candidate priming sites are defined as primer-length segments of the consensus whose summed entropy does not exceed a cutoff (div_cut; default: 2.0). Segments are paired when the difference between their forward and reverse primer start positions falls within the target amplicon-size range (default: 300–1000 bp) and their GC difference is below tolerance (GC_tol; default: 0.10), with the upstream segment taken as the forward primer and the reverse complement of the downstream segment as the reverse primer. Ranked candidates are subsequently screened for secondary structure, and pairs predicted to form stable hairpins or self-dimers are discarded. Because conserved priming sites tend to flank variable regions, and because the input alignment is restricted to a phylogenetically informative gene, the resulting amplicons capture the sequence diversity required for species-level discrimination.

The ten highest-ranked candidate pairs that pass secondary-structure screening are validated *in silico* against every downloaded assembly using *seqkit locate* [19]. Both strands are scanned, and each primer is matched as an IUPAC-degenerate pattern, so that every sequence variant encoded by a degenerate position is treated as an exact match; no additional mismatches are permitted. All same-strand combinations of forward and reverse binding sites falling within the permitted amplicon-size range are reconstructed, and the candidate with the best validation performance is recommended, which may therefore differ from the pair ranked highest at the design stage. Product lengths reported at this stage include both primer-binding segments, whereas design-stage spacing is measured between primer start positions. For each gene, the pipeline reports the number and proportion of genomes successfully amplified, the mean amplicon length, and species-level metrics comprising the number of amplified species, the proportion of amplified genomes carrying multiple intragenomic amplicon alleles, and the proportion of species showing inter-species allele overlap. Results are compiled into an HTML report; when multiple genes are analyzed, an additional cross-gene comparison report is generated.

### Implementation

The workflow environment is pinned with Pixi, which fixes Python, NumPy, and the command-line tools VSEARCH, MUSCLE, and SeqKit across Linux, macOS, and Windows. A dependency-resolution layer detects the required external tools at run time and installs missing executables where possible. Provenance is recorded at every step: genome downloads are fingerprinted with SHA-256 checksums, the active configuration is hashed, and a download manifest ties each run to its inputs. Correctness is guarded by an automated functional-test suite, a performance smoke test, and continuous integration on all three platforms. The workflow is exposed as both a command-line interface and a lightweight Python API (AmPairProject), making it embeddable in larger analyses.

## Results

### Species-level resolution and community application of the designed *Bacillus* primer set

Using the genus *Bacillus* as a case study, we selected *tuf* as the candidate gene. This gene encodes the elongation factor thermal unstable (EF-Tu) and is a frequently reported discriminatory housekeeping marker for *Bacillus*. We then retrieved 2,392 complete genomes from NCBI. After dereplication and multiple-sequence alignment, we identified conserved, low-entropy priming sites at alignment positions 58, 517 and 1,004.

Among the primer pairs formed from these sites, the highest-scoring combination placed the forward primer at position 58 and the reverse primer near position 1,004, spanning a 965-bp variable region that enables species-specific identification. This combination was ranked highest by low priming-site entropy (sequence diversity) and small GC-content difference, as described in Methods. The resulting pair (forward, 5′-CACGTTGACCAYGGTAAAACH-3′; reverse, 5′-GTDAYRTCHGWWGTACGGA-3′) amplified 99% of the downloaded genomes *in silico*.

To evaluate the improved discrimination power of the *tuf* primer sets, we used RibDif2 to assess the metataxonomic performance of each primer set *in silico* on the 2,392 genomes downloaded from NCBI (July 2026) [14]. The *tuf* primer set amplified 2,390 genomes, of which only 1 (0.04%) carried multiple alleles, and none of the 59 amplified species (0.0%) showed any inter-species allele overlap. By contrast, the 16S rRNA V1– V9 primers (forward, 5′-AGRGTTYGATYMTGGCTCAG-3′; reverse, 5′-RGYTACCTTGTTACGACTT-3′) amplified 2,375 genomes, among which 2,171 (91.41%) carried multiple alleles, and 41 of 59 (69.49%) amplified species exhibited at least one inter-species allele overlap. Consistent with these results, the RibDif2 network visualization of genome overlap showed that amplicons generated by the designed *tuf* primer set fully resolved individual *Bacillus* species (Fig. 3A), whereas the 16S rRNA amplicons produced extensive allele overlap among species (Fig. 3B). Together, these results demonstrate that the primer set recommended by AmPair provide species-level discrimination power for amplicon sequencing.

**Figure 3.**
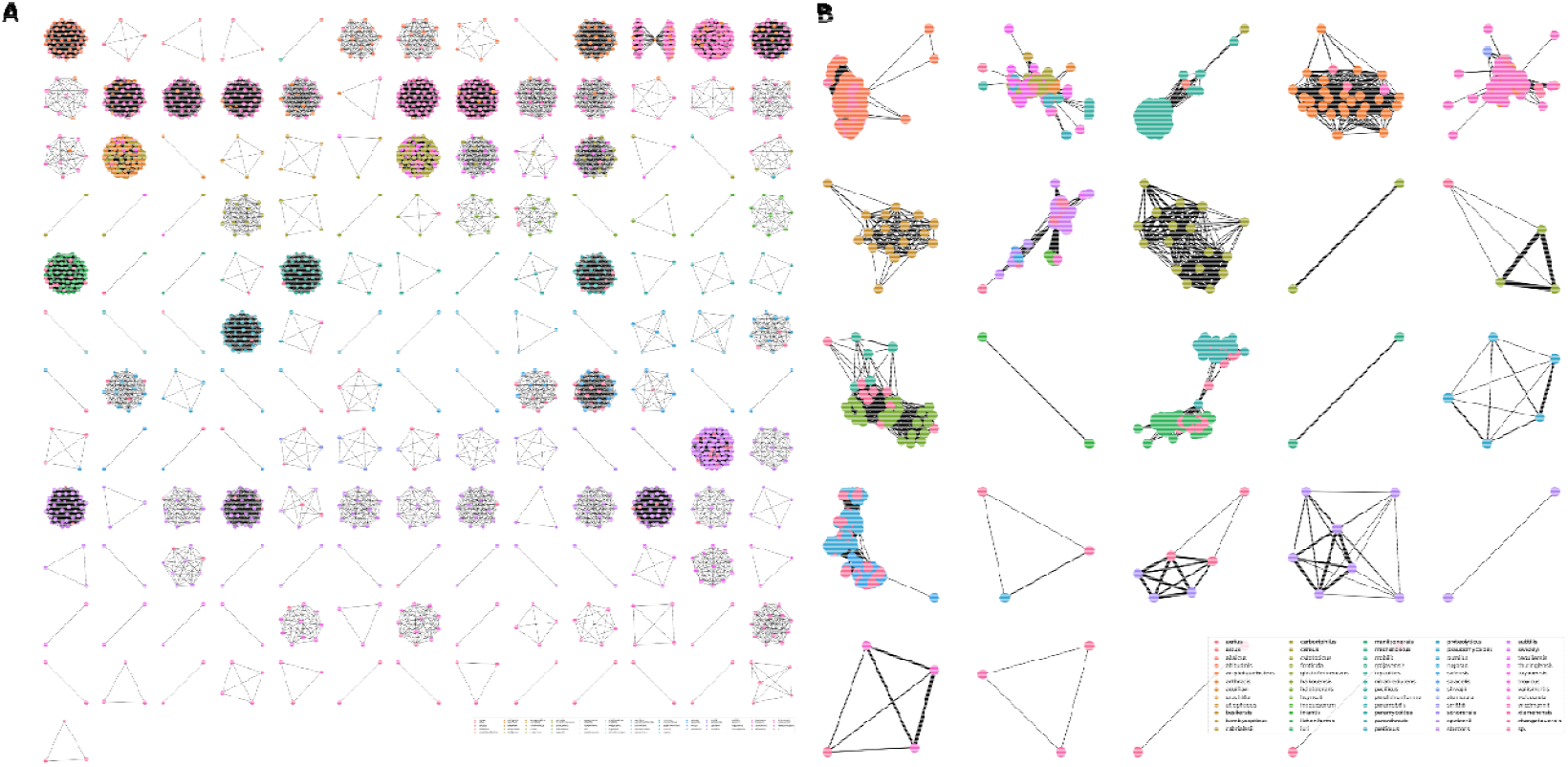
Network visualization of *Bacillus* genome overlaps generated by RibDif2. (A) Genome overlaps based on *tuf* amplicons. (B) Genome overlaps based on the 16S rRNA V1–V9 region. Each node represents a *Bacillus* genome, and two nodes are connected when their genomes share overlapping alleles. Node color denotes the *Bacillus* species. Edge width is proportional to the number of shared alleles. Non-connected nodes are excluded.

To demonstrate the practical application of the *tuf* primer set beyond *in silico* benchmarking, we applied the primer to a lignin-enriched biofilm assembled from a *Bacillaceae* strain library. Nanopore-based metataxonomic profiling with these primers resolved the community at the species level, distinguishing closely related *Bacillus* taxa, including *Bacillus paralicheniformis, Bacillus subtilis, Bacillus velezensis*, and *Bacillus licheniformis*. This confirms that the primer set recommended by AmPair are applicable to metataxonomic studies of complex communities, not only to benchmark genome datasets.

## Discussion

In this study, we developed AmPair, a framework for recommending and designing housekeeping-gene primer set that enable species-level resolution in amplicon sequencing. Using *Bacillus* as a demonstration, we showed that the *tuf*-targeted primers recommended by AmPair discriminate closely related species that standard 16S rRNA amplicon sequencing cannot resolve, and we validated this both *in silico* across 2,392 genomes and in a *Bacillaceae* synthetic microbial community.

This improved resolution reflects fundamental differences between the two markers. The single-copy, more rapidly evolving housekeeping gene largely eliminates the artifacts arising from the multiple alleles and inter-species overlap that affect 16S rRNA, yielding amplicons that resolve individual species. Beyond this gain in resolution, housekeeping-gene amplicon sequencing occupies a cost-effective middle ground between shotgun metagenomics and full-length 16S rRNA sequencing. Rather than relying on a single fixed marker, AmPair tailors primer selection to the target taxa and to the most informative gene, making species-level amplicon profiling broadly accessible. Several limitations should be noted. First, AmPair depends on the availability and quality of reference genomes, and taxa that are underrepresented in public databases may yield less reliable primer recommendations. Second, primer performance was assessed largely *in silico*; although the *tuf* primers have since been applied experimentally, *in silico* amplification does not fully capture PCR efficiency, chimera formation, or other experimental biases. We therefore recommend that users carefully evaluate their choice of housekeeping gene and experimentally validate the primer set suggested by AmPair. Housekeeping-gene amplicon sequencing is increasingly recognized as a powerful complement to 16S rRNA profiling. Single-copy protein-coding markers such as *rpoD, gyrA*, and *gyrB* evolve more rapidly and lack the intragenomic allelic heterogeneity of the rRNA operon, and have been used to achieve species- or even strain-level resolution across diverse bacterial groups [15,20–23]. Their applications range from tracking pathogens in food and clinical samples to dissecting closely related taxa in complex environmental communities, where the limited resolution of 16S rRNA often leaves species indistinguishable. Broader adoption has been limited mostly by the effort required to identify a suitable marker and design discriminating primers for each target taxon. AmPair addresses this bottleneck by automating marker-aware primer design, extending readily to a range of housekeeping genes across diverse genera. Thus, AmPair offers a generalizable route to species-level community profiling that is accurate, accessible, and readily deployable in real-world applications.

Beyond the biological contribution, AmPair also addresses the practical barriers that have slowed the adoption of such pipelines. A pinned, cross-platform environment, automated dependency resolution, and built-in provenance tracking lower the technical threshold for non-expert users, while the functional and performance test suites and continuous integration across three operating systems provide the reproducibility expected of a tool intended for routine use. By exposing both a command-line interface and a Python API, AmPair can be embedded into larger metataxonomic workflows rather than remaining a single-use script. These engineering choices, although secondary to the primer-design methodology, are what make the method broadly deployable in practice.

## Acknowledgements

X.X. was supported by start-up funding from the Institute of Biology Leiden, Leiden University, awarded to Á.T.K. The authors thank members of the Nadine Ziemert lab for helpful discussions during X.X.’s research visit to the University of Tübingen.

## References

1. Lundberg, D.S. et al. (2013) Practical innovations for high-throughput amplicon sequencing. Nat. Methods 10, 999–1002

2. Kozich, J.J. et al. (2013) Development of a dual-index sequencing strategy and curation pipeline for analyzing amplicon sequence data on the MiSeq illumina sequencing platform. Appl. Environ. Microbiol. 79, 5112–5120

3. Schirmer, M. et al. (2015) Insight into biases and sequencing errors for amplicon sequencing with the illumina MiSeq platform. Nucleic Acids Res. 43, e37

4. Schoch, C.L. et al. (2012) Nuclear ribosomal internal transcribed spacer (ITS) region as a universal DNA barcode marker for fungi. Proc. Natl. Acad. Sci. 109, 6241–6246

5. Větrovský, T. and Baldrian, P. (2013) The variability of the 16S rRNA gene in bacterial genomes and its consequences for bacterial community analyses. PLOS ONE 8, e57923

6. Barb, J.J. et al. (2016) Development of an analysis pipeline characterizing multiple hypervariable regions of 16S rRNA using mock samples. PLOS ONE 11, e0148047

7. Rhoads, A. and Au, K.F. (2015) PacBio sequencing and its applications. Genomics Proteomics Bioinformatics 13, 278–289

8. Brown, C.G. and Clarke, J. (2016) Nanopore development at oxford nanopore. Nat. Biotechnol. 34, 810–811

9. Bertolo, A. et al. (2024) Optimized bacterial community characterization through full-length 16S rRNA gene sequencing utilizing MinION nanopore technology. BMC Microbiol. 24, 58

10. Matsuo, Y. et al. (2021) Full-length 16S rRNA gene amplicon analysis of human gut microbiota using MinIONTM nanopore sequencing confers species-level resolution. BMC Microbiol. 21, 35

11. Johnson, J.S. et al. (2019) Evaluation of 16S rRNA gene sequencing for species and strain-level microbiome analysis. Nat. Commun. 10, 5029

12. Xu, X. et al. (2023) Enhanced specificity of Bacillus metataxonomics using a tuf-targeted amplicon sequencing approach. ISME Commun. 3, 1–11

13. Strube, M.L. (2021) RibDif: Can individual species be differentiated by 16S sequencing? Bioinforma. Adv. 1, vbab020

14. Murphy, R. and Strube, M.L. (2023) RibDif2: Expanding amplicon analysis to full genomes. Bioinforma. Adv. 3, vbad111

15. Lauritsen, J.G. et al. (2021) Identification and Differentiation of Pseudomonas Species in Field Samples Using an rpoD Amplicon Sequencing Methodology. mSystems 6, 10.1128/msystems.00704-21

16. Strube, M.L. et al. (2018) A detailed investigation of the porcine skin and nose microbiome using universal and Staphylococcus specific primers. Sci. Rep. 8, 12751

17. Rognes, T. et al. (2016) VSEARCH: a versatile open source tool for metagenomics. PeerJ 4, e2584

18. Edgar, R.C. (2022) Muscle5: High-accuracy alignment ensembles enable unbiased assessments of sequence homology and phylogeny. Nat. Commun. 13, 6968

19. Shen, W. et al. (2016) SeqKit: A cross-platform and ultrafast toolkit for FASTA/Q file manipulation. PLOS ONE 11, e0163962

20. Ren, Q. and Hill, J.E. (2023) Rapid and accurate taxonomic classification of cpn60 amplicon sequence variants. ISME Commun. 3, 1–7

21. Bisaschi, M. et al. (2025) Nanopore-based amplicon sequencing for rapid detection and identification of bacillus spp. in plant-based products. Front. Microbiol. 16

22. Liu, Y. et al. (2022) Housekeeping gene gyrA, a potential molecular marker for Bacillus ecology study. AMB Express 12, 133

23. Poirier, S. et al. (2018) Deciphering intra-species bacterial diversity of meat and seafood spoilage microbiota using gyrB amplicon sequencing: A comparative analysis with 16S rDNA V3-V4 amplicon sequencing. PLOS ONE 13, e0204629

